# Cylindromatosis Drives Synapse Pruning and Weakening by Promoting Macroautophagy through Akt-mTOR Signaling

**DOI:** 10.1101/2021.12.08.471792

**Authors:** Alexis S. Zajicek, Hongyu Ruan, Huihui Dai, Mary C. Skolfield, Hannah L. Phillips, Wendi J. Burnette, Behnam Javidfar, Shao-Cong Sun, Schahram Akbarian, Wei-Dong Yao

## Abstract

The lysine-63 deubiquitinase cylindromatosis (CYLD) is long recognized as a tumor suppressor in immunity and inflammation and its loss-of-function mutations lead to familial cylindromatosis. However, recent studies reveal that CYLD is enriched in mammalian brain postsynaptic densities, and a gain-of-function mutation causes frontotemporal dementia (FTD), suggesting critical roles at excitatory synapses. Here we report that CYLD drives synapse elimination and weakening by acting on the Akt-mTOR-autophagy axis. Mice lacking CYLD display abnormal sociability, anxiety- and depression-like behaviors, and cognitive inflexibility. These behavioral impairments are accompanied by excessive synapse numbers, increased postsynaptic efficacy, augmented synaptic summation, and impaired NMDA receptor-dependent hippocampal long-term depression (LTD). Exogenous expression of CYLD results in removal of established dendritic spines from mature neurons in a deubiquitinase activity-dependent manner. In search of underlying molecular mechanisms, we find that CYLD knockout mice display marked overactivation of Akt and mTOR and reduced autophagic flux and, conversely, CYLD overexpression potently suppresses Akt and mTOR activity and promotes autophagy. Consequently, abrogating the Akt-mTOR-autophagy signaling pathway abolishes CYLD-induced spine loss, whereas enhancing autophagy in vivo by the mTOR inhibitor rapamycin rescues the synaptic pruning and LTD deficits in mutant mice. Our findings establish CYLD, via Akt-mTOR signaling, as a synaptic autophagy activator that exerts critical modulations on synapse maintenance, function, and plasticity.

## INTRODUCTION

The tumor suppressor *CYLD* encodes a lysine-63 (K63) deubiquitinating enzyme (DUB) [1] and its loss-of-function mutations cause familial cylindromatosis, an autosomal dominant predisposition to head and neck skin tumors [2]. Unlike lysine-48 (K48)-linked polyubiquitin chains that tag protein substrates for proteasomal degradation, K63 chains facilitate multiprotein complex assembly during signaling transduction, and are classically involved in the NF-κB pathway that mediates immune response and inflammation [3-5]. Landmark studies have established that CYLD suppresses tumorigenesis and promotes apoptosis by cleaving K63 chains on key components of the NF-kB pathway, inhibiting NF-κB activation [6-8]. CYLD also negatively regulates other processes, such as TGFβ signaling via deubiquitinating its substrate Akt/PKB [9]. Mice deficient in CYLD display modest immune phenotypes as they do not develop tumors spontaneously but show increased susceptibility to tumorigenesis and other immune deficits when challenged [10-12].

CYLD is rapidly emerging as a major player at synapses in the mammalian brain [13]. CYLD is enriched in the postsynaptic density (PSD) of excitatory synapses [14, 15], is recruited to the PSD in response to activity [16], and interacts with >100 synaptic proteins in mouse striatal synaptosomes [17]. CYLD deubiquitinates and regulates synaptic abundance of the postsynaptic scaffold PSD-95, regulates synaptic strength, and mediates NMDA receptor (NMDAR)-dependent chemical LTD (cLTD) in cultured neurons [15]. Although most heavily enriched in the PSD, CYLD is also present in the presynaptic fraction of rodent brain [15] and regulates axonal length in cultured mouse hippocampal neurons [18]. In addition, CYLD has a role in inhibitory synaptic transmission and regulates GABA receptor trafficking/turnover in striatal neurons [19]. Finally, a gain-of-function mutation and variants of *CYLD* were identified in patients with frontotemporal dementia (FTD)/amyotrophic lateral sclerosis (ALS) [18, 20], placing *CYLD* as the newest member of the FTD/ALS gene family. However, the precise role of CYLD in synaptic remodeling, plasticity, and behavior *in vivo* remains undetermined.

Macroautophagy (autophagy) is a conserved catabolic process that delivers unnecessary or damaged cytosolic constituents to lysosomes for degradation to maintain cellular homeostasis [21, 22]. Autophagy is particularly important in neurons, which are postmitotic and must thrive throughout an organism’s lifetime. Indeed, autophagy plays essential roles in neuron viability and development [23, 24] and is implicated in numerous neurodegenerative and psychiatric diseases [25-29]. Increasing evidence also supports critical roles for neuronal autophagy in synaptic function, remodeling, and plasticity [30-32]. In particular, neuronal autophagy is induced during, and required for, NMDAR-dependent LTD [33-35], and autophagy induction promotes hippocampal long-term potentiation (LTP) and memory and reverses age-related memory decline [36]. Furthermore, neuronal autophagy promotes synapse pruning in human and mouse cortex during adolescent development [37]. mTOR, a multipurpose serine/threonine kinase localized in the synapse, is a central regulator of autophagy. Together with its upstream activator Akt, mTOR inhibits autophagosome formation, the induction step of autophagy [38, 39]. Impaired mTOR-autophagy is involved in aberrant synaptic pruning associated with autism [37] and synaptic and cognitive deficits in a mouse model of fragile X syndrome [40].

In this study, we show that CYLD drives synapse pruning, weakens synaptic strength, and mediates LTD by stimulating Akt-mTOR-autophagy signaling, and is required for maintenance of social, affective, and cognitive behaviors.

## RESULTS

### Aberrant social, affective, and cognitive behaviors in CYLD-deficient mice

To gain initial insights into in vivo roles of CYLD in the brain, we subjected CYLD WT and KO mice to a battery of behavioral tests (Figure 1). CYLD KO mice travelled significantly less distance and spent less time in the center of an open field compared to WT littermates (Figure 1A), suggesting impaired locomotor activity and/or reduced motivation to explore, as well as increased anxiety-like behavior. KO mice displayed longer latency and fewer crossings to light in a Light/Dark box (Figure 1B) and spent more time in the closed arms and had fewer entries to the open arms in an elevated plus maze (Figure 1C), signaling heightened anxiety. KO mice showed significant increases in the duration of immobility than WT in both forced swim and tail suspension tests (Figure 1D and 1E), suggesting a depression phenotype. CYLD KO mice spent significantly more time interacting with a stranger conspecific compared to WT, though the time in each chamber remained unchanged, suggesting increased sociability (Figure 1F). Working memory appeared unaltered in KO mice as they showed similar spontaneous alternations to WT in a Y-maze (Figure S1). Finally, when placed in a Barnes maze, KO mice performed equally well as WT when learning to find an escape tunnel, but had difficulty in re-learning the escape tunnel when it was in the opposite location (Figure 1G and 1H), suggesting impaired reversal learning and cognitive flexibility.

**Figure 1.**
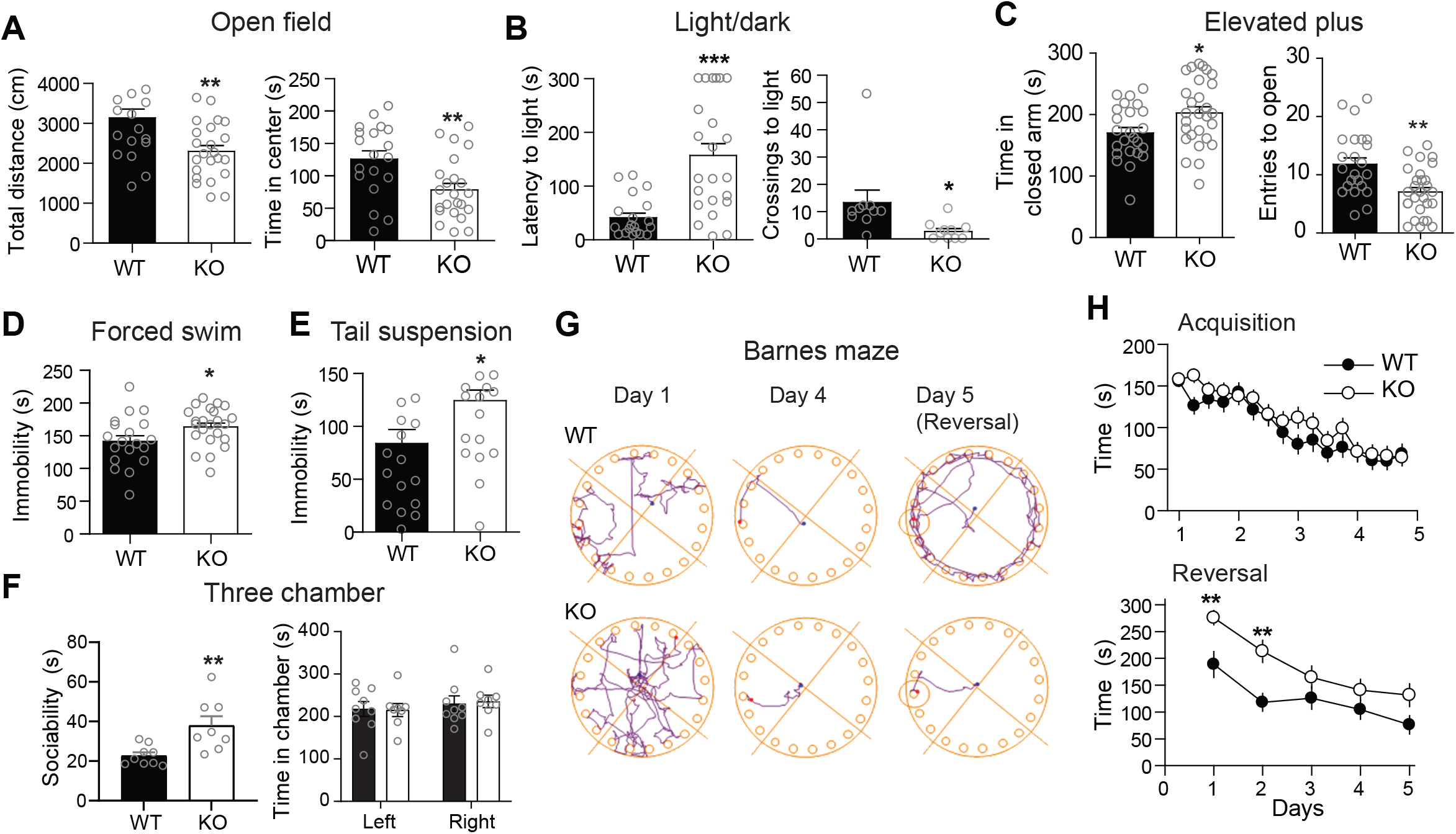
Aberrant locomotor, social, affective, and cognitive behaviors in CYLD KO mice. (A) Open field test. (B) Light/Dark test. (C) Elevated plus maze test. (D) Forced-swim test. (E) Tail-suspension test. (F) Three-chamber sociability test. (G) Representative track plots from mice during a Barnes maze test. Blue dot indicates starting point, red dot indicates end point. On reversal day 5, the circle represents the area around the previous escape tunnel. (H) Barnes maze quantifications. Time represents latency to escape during a 3 minute test four times daily (top) or a 5 minute single trial each day during reversal (bottom). n = 19 (WT) and 23 (KO) mice for (A-E); n = 8 (WT) and 9 (KO) mice for (F); n = 30 (WT) and 33 (KO) mice for G,H). *p < 0.05; **p < 0.01; ***p < 0.001. Two-tailed unpaired t-tests. n = 30 (WT) and 33 (KO) mice. Summary data in all figures are mean ± sem.

### Enhanced postsynaptic efficacy in CYLD KO mice

We next performed whole-cell patch-clamp analyses on hippocampal CA1 and medial prefrontal cortex (mPFC) layer 5 pyramidal neurons in slices prepared from WT and KO mice (Figure 2). In CA1 (P20-P30), the frequency of AMPA receptor (AMPAR)-mediated miniature excitatory postsynaptic currents (mEPSCs) was significantly higher in KO neurons compared to WT with the mEPSC amplitude being indistinguishable between the two groups (Figure 2A and 2B). In mPFC, mEPSC frequency was also significantly higher in P60 KO neurons compared to WT (Figure S2A-C). These results suggest an increase in presynaptic glutamate release probability and/or number of functional synapses in mutant neurons.

**Figure 2.**
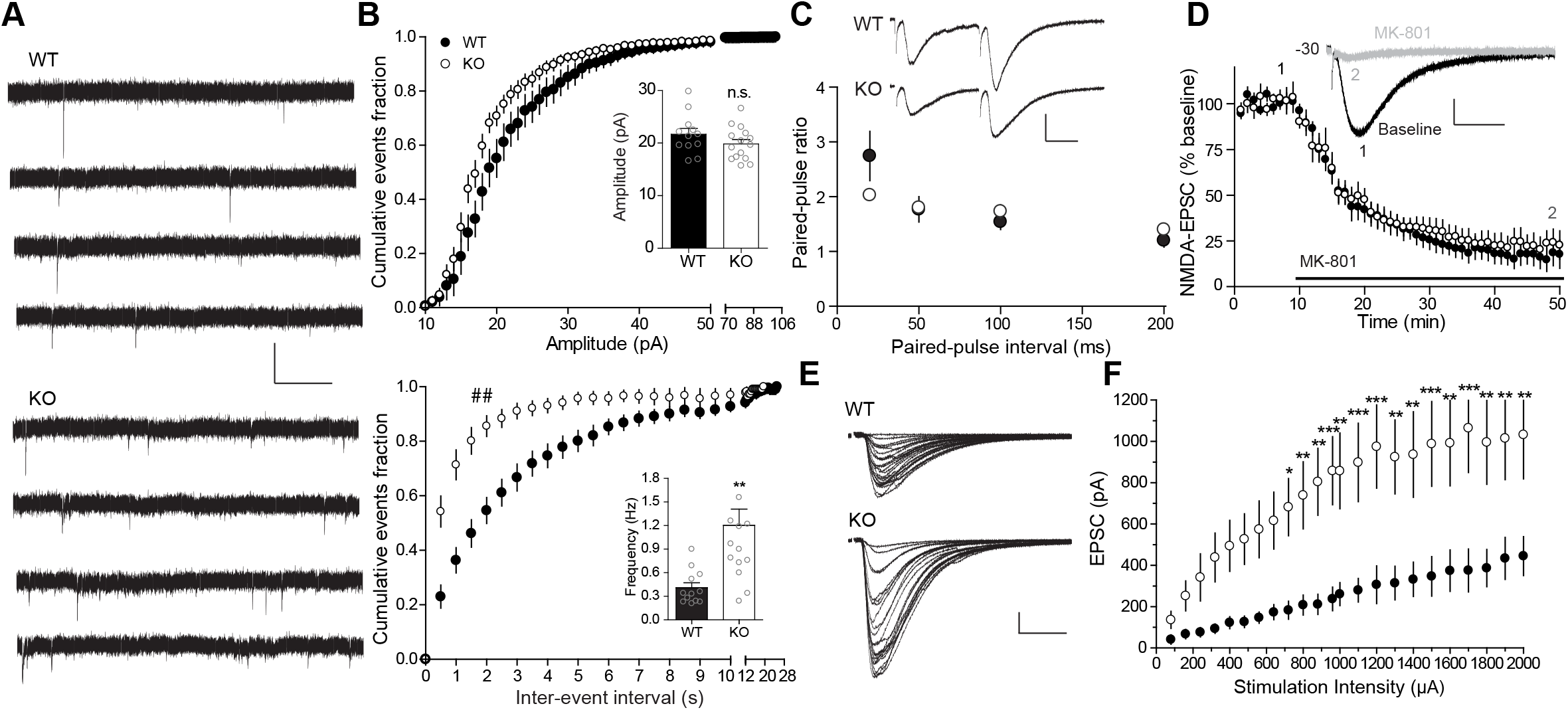
Enhanced postsynaptic efficacy in CYLD KO mice. (A) Representative mEPSC traces from CA1 hippocampal neurons. Scale bar: 40 pA, 2 sec. (B) Cumulative probability distributions of mEPSC amplitude and inter-event intervals. Insets, Mean amplitudes and frequencies. ## p < 0.01, Kolmogorov–Smirnov test; **p < 0.01; n.s., not significant; two-tailed unpaired t tests. (C) Paired-pulse ratio. Scale bar:100 pA, 20 ms. n = 8 (WT) and 8 (KO) cells. (D) MK-801 assay. NMDA-EPSCs before and after MK-801 (20 μM) wash-in were recorded at -30 mV. Inset, representative NMDA-EPSC before (1) and after (2) MK-801. Scale bar: 100 pA, 50 ms. n = 6 (WT) and 6 (KO) cells. (E,F) Representative (E) and summary (F) EPSCs in response to varying stimulation intensities at Schaffer collateral-CA1 synapses. Scale bar: 200 pA, 20 ms. n = 16 (WT) and 10 (KO) cells. * p < 0.05, ** p < 0.01, *** p < 0.001, Two-way ANOVA with Bonferroni’s multiple comparisons test.

To further delineate the underlying mechanisms, we examined the evoked EPSCs of Shaffer collateral-CA1 synapses. The paired-pulse ratios (PPRs), a measure of changes in presynaptic release probability [41], were not significantly different at all intervals tested between KO and WT neurons (Figure 2C), suggesting unaltered presynaptic mechanisms in mutant neurons. To further verify this, we performed a use-dependent NMDAR blockade assay with MK-801, an open-channel blocker of NMDARs. MK-801 more rapidly inhibits NMDARs at synapses with a high release probability, due to more frequent opening of these receptors, than those at low-release probability synapses. MK-801 had similar NMDAR-EPSC inhibition time courses in CYLD KO and WT neurons (Figure 2D), confirming an intact glutamate release mechanism in mutant neurons. Finally, we examined the input-output (I-O) relationship of AMPAR-EPSCs in response to increasing presynaptic stimulation intensities. EPSC amplitudes were markedly higher across all stimulus intensities in CYLD KO neurons compared to WT (Figure 2E and 2F). Together, these results demonstrate that synaptic strength in CYLD KO mice is significantly increased, likely due to more functional synapses in mutant mice.

### Augmented short-term but impaired long-term synaptic plasticity in CYLD KO mice

We next examined short-and long-term synaptic plasticity of Shaffer collateral-CA1 synapses (Figure 3). Synaptic responses under current-clamp to high-frequency train stimulation allow assessment of summation and sustainment of postsynaptic depolarization at physiological firing frequencies [42]. WT neurons displayed a characteristic summation of postsynaptic depolarization in response to 15 20-Hz stimuli, which was significantly increased in CYLD KO neurons, suggesting enhanced short-term plasticity (Figure 3A-3B). Under voltage-clamp, a conventional protocol involving pairing low-frequency (2 Hz) presynaptic stimulation with postsynaptic depolarization produced robust NMDAR-dependent LTP in WT (147.8 ± 13.4%); this LTP displayed a trend toward impairment in CYLD KO neurons (117.6 ± 15.8%; P = 0.18 vs. WT; t-test) (Figure 3C). Furthermore, NMDAR-dependent LTD induced by a standard low-frequency stimulation protocol (1000 pulses at 2 Hz) in WT (69.759 ± 5.6%) was absent in CYLD KO neurons (91.6 ± 9.0%; P = 0.04 vs. WT; t-test). In contrast, mGluR1-dependent LTD induced by the group I metabotropic glutamate receptor (mGluR1) agonist DHPG was similar in WT and KO neurons (Figure 3E; P = 0.47 vs. WT; t-test). Together, these results demonstrate that CYLD plays important roles in maintaining both short-term and NMDAR-dependent long-term synaptic plasticity.

**Figure 3.**
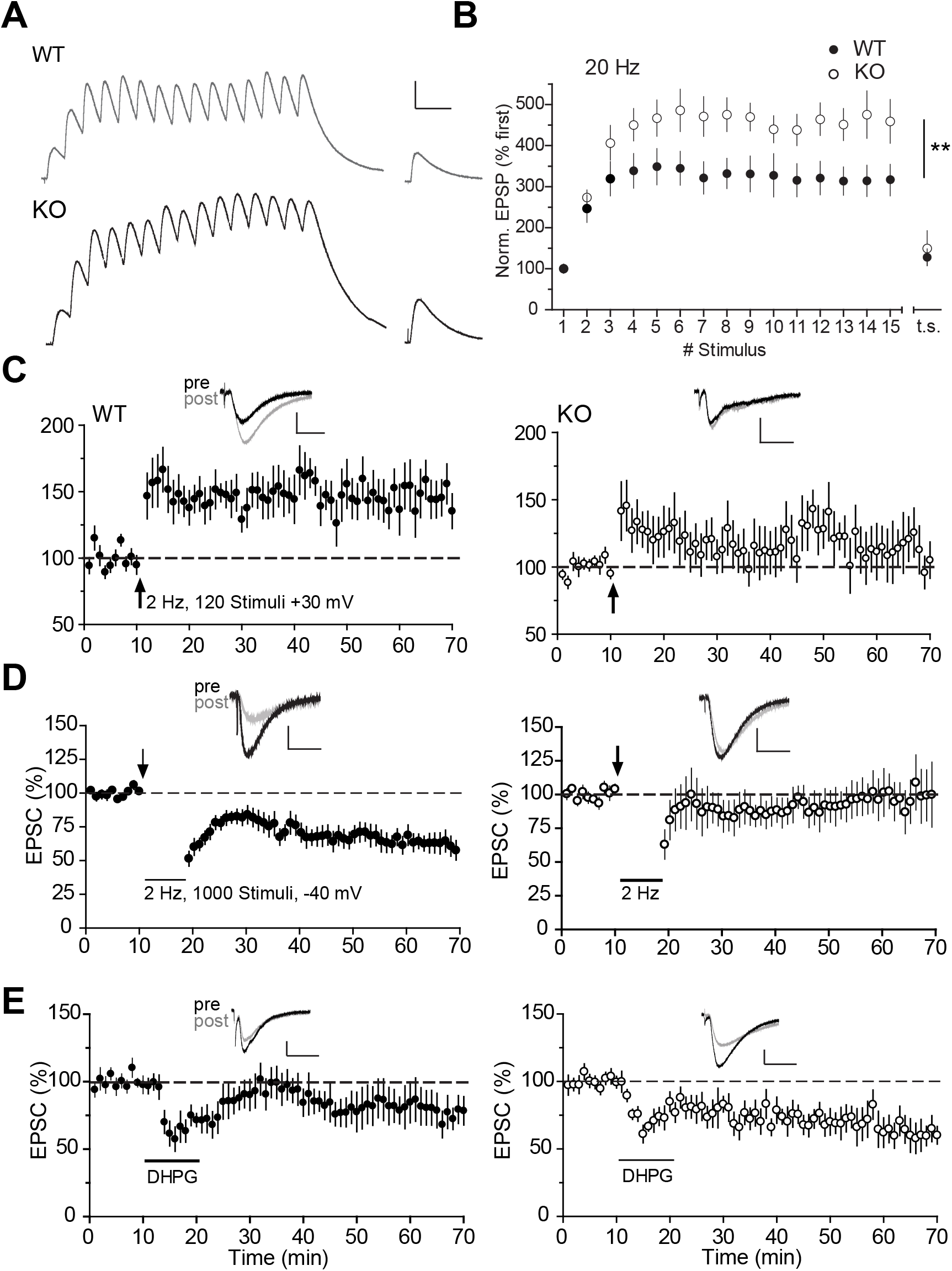
Augmented short-term but impaired long-term synaptic plasticity in CYLD KO mice. (A). Synaptic summation as shown by representative EPSPs in response to a 15-pulse stimulus train at 20 Hz followed by a recovery test pulse. Scale bar: 5 mV, 100 ms. (B) Quantifications of synaptic summation from (A), shown as summary of normalized EPSP amplitudes (to the peak of first EPSP) versus stimulus sequence during the train. n = 9 (WT) and 8 (KO) cells. ** p < 0.01, Two-way ANOVA. (C) LTP, induced (arrows) by 120 stimuli at 2 Hz delivered to the Schaffer collateral afferent while holding CA1 cells at +30 mV. Scale bars: 50 pA, 25 ms. n = 8 (WT) and 10 (KO) cells. (D) LTD, induced (arrows) by 1000 stimuli at 2 Hz delivered to the Schaffer collateral afferent while holding CA1 cells at -40 mV. Scale bars: 50 pA, 25 ms. n = 12 (WT) and 7 (KO) cells. (E) mGluR1-LTD. DHPG (75 μM) was washed in between 10-20 min. n = 7 (WT) and 6 (KO) cells. Insets, representative EPSCs before (pre) and after (post) respective LTP/LTD induction procedures. Scale bars: 50 pA, 25 ms. All LTP/LTD sample traces are averages of 5 EPSCs.

### CYLD promotes synapse pruning in a DUB-dependent manner

We next investigated synaptic alterations in CYLD KO mice at biochemical and structural levels and the roles of CYLD in dendritic spine remodeling (Figure 4). Immunoblotting (IB) analysis revealed significantly higher levels of hippocampal GluA1, GluN1, and PSD-95 in CYLD KO mice (Figure 4A and 4B), consistent with increased postsynaptic efficacy. We next compared dendritic spine density of GFP-labeled neurons in P30 WT and KO mice by imaging CA1 hippocampal neurons injected on P15 with AAV2-hSyn-EGFP. KO neurons had a significant increase in numbers of total and mushroom (believed to be associated with mature, stable synapses[43]) spines (Figure 4C and 4D). These data demonstrate that CYLD KO neurons possess excessive numbers of synapses.

**Figure 4.**
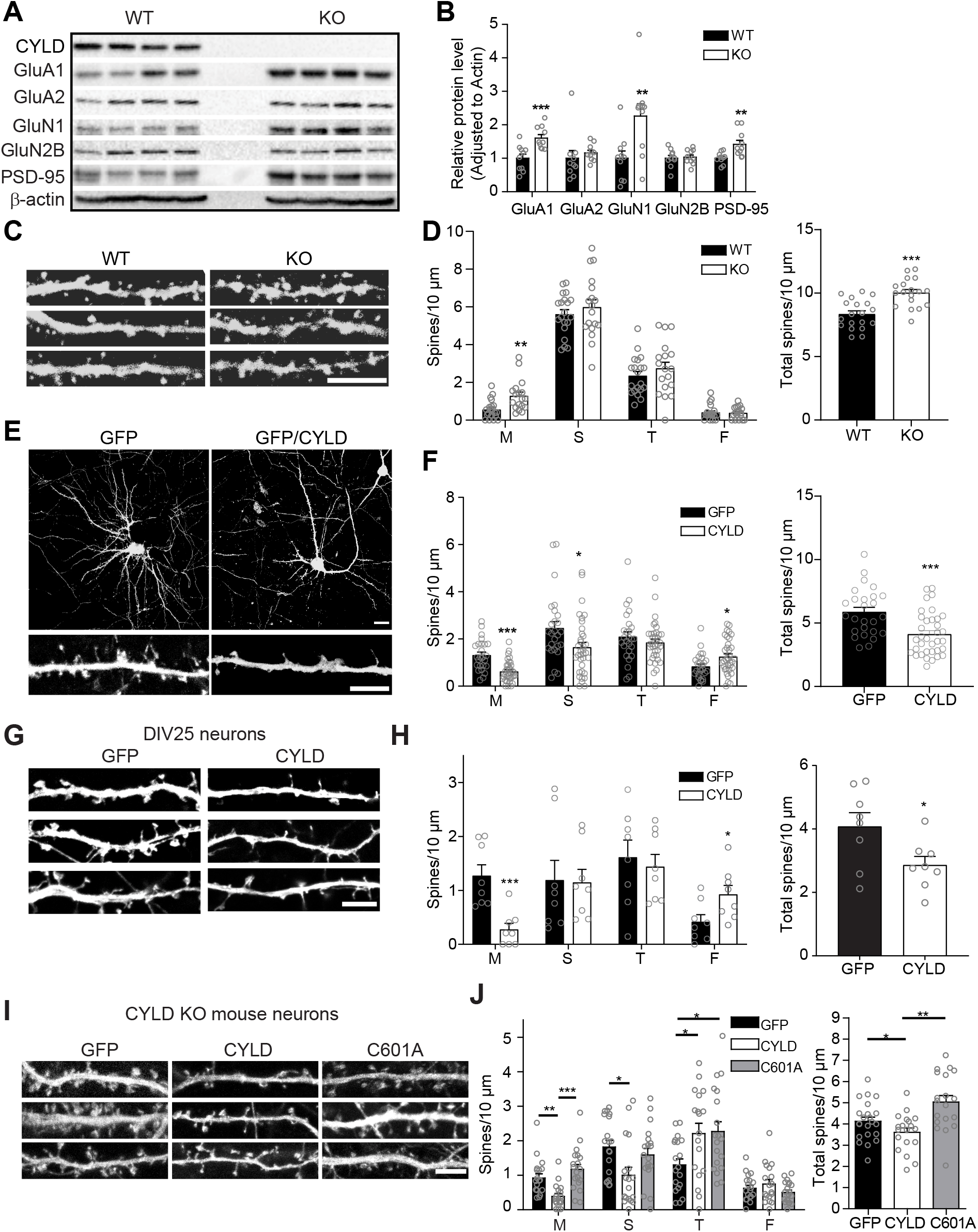
CYLD promotes synapse pruning. (A) Representative IBs of indicated synaptic proteins on CYLD WT and KO hippocampus lysates. (B) Quantification of protein levels from (A) normalized to WT controls. n = 10 (WT) and 10 (KO) mice. (C) Representative images of dendritic segments from AAV2-hSyn-EGFP infected CA1 hippocampal neurons. Scale bar: 5 μm. (D) Quantifications of dendritic spine density from (C). n = 20 (WT) and 18 (KO) cells from 3 mice in each group. (E) Representative images of whole neurons (top) and dendritic segments (bottom) (DIV21) from GFP or GFP/CYLD transfected (DIV7) rat embryonic hippocampal neurons. Scale bars: 20 μm (top), 5 μm (bottom). (F) Quantifications of dendritic spine densities from (E). M, mushroom; S, stubby; T, thin; F, filopodia. n = 26 (GFP) and 35 (GFP/CYLD) cells from 3-4 independent cultures. (G, H) Effects of CYLD overexpression in mature neurons. Representative dendritic segments (G) and quantifications (H) from mature rat hippocampal neurons transfected with GFP or GFP/CYLD on DIV19 and imaged on DIV25. Scale bar: 5 μm. n = 8 (GFP) and 10 (GFP/CYLD) from 3 independent experiments. (I, J). Effects of CYLD and C601A in spine densities in cultured CYLD KO mouse neurons. Scale bar: 5 μm. n = 21 (GFP), 19 (CYLD), and 19 (C601A). *p < 0.05; **p < 0.01; ***P < 0.001; two-tailed unpaired t-tests or one-way ANOVA with post-hoc Bonferroni’s multiple comparison test.

To further investigate the causal role of CYLD in spine remodeling, we examined if CYLD overexpression can drive dendritic spine removal. Exogenous expression of CYLD in cultured rat hippocampal neurons resulted in significantly decreased total, mushroom, and stubby spine densities paralleled by an increase in filopodia protrusions (Figure 4E and 4F), as well as significantly decreased GluA1 and GluN1 levels (Figure S2). To distinguish whether CYLD-mediated spine removal was due to defective spine formation or facilitated spine pruning, we repeated above experiments in older neurons (Figure 4G and 4H). Transfecting CYLD into neurons at DIV19, when the majority of synapses have been established and matured, still resulted in significant elimination of total and mushroom spines correlated with an increase of filipodia protrusions (Figure 4G and 4H). Finally, to test if CYLD modulation of spines depended on its DUB activity, we expressed GFP control, wild-type CYLD, or a DUB-deficient CYLD (C601A) mutant in cultured CYLD KO hippocampal neurons where endogenous CYLD was absent to eliminate potential confounding effects. We found that wild-type CYLD, but not the enzyme-dead C601A, significantly reduced total, mushroom, and stubby spine densities (Figure 4I and 4J). Together, these results demonstrate that CYLD, in a DUB-dependent manner, drives pruning of existing glutamatergic synapses on mature neurons.

### Impaired autophagy and hyperactive Akt-mTOR signaling in CYLD KO brains

We next investigated the mechanism by which CYLD drives synapse removal. We focused on autophagy because of its roles in synapse pruning [37, 40, 44] and the potential links between CYLD and autophagy via K63 ubiquitination-related mechanisms [45-47]. There were significantly lower levels of LC3-II, the lipidated form of LC3-I and the only reliable marker for autophagosomes, and LC3-II/I ratio in KO hippocampal lysates at P30, P60, and P270 compared to WT (Figure 5A and 5B). In addition, there was a significantly higher level of p62, an autophagy substrate and a functional readout of lysosomal degradation known to accumulate when autophagy is compromised, at all ages (Figure 5A and 5B), supporting reduced autophagic flux in KO mice. LC3-I and LC3-II were also both significantly decreased in CYLD KO cortex (Figure S4A and 4B). LC3 immunofluorescence showed significant reduction of LC3 puncta numbers in somas of CA1 neurons on hippocampal sections from KO mice, confirming reduced autophagosome formation (Figure 5C and 5D). Finally, CYLD overexpression in cultured hippocampal neurons resulted in a significant increase in LC3-II (Figure 5C and 5D), confirming that reduced autophagy in KO was due to a loss of CYLD. Together, these results demonstrate that CYLD promotes autophagy in neurons.

**Figure 5.**
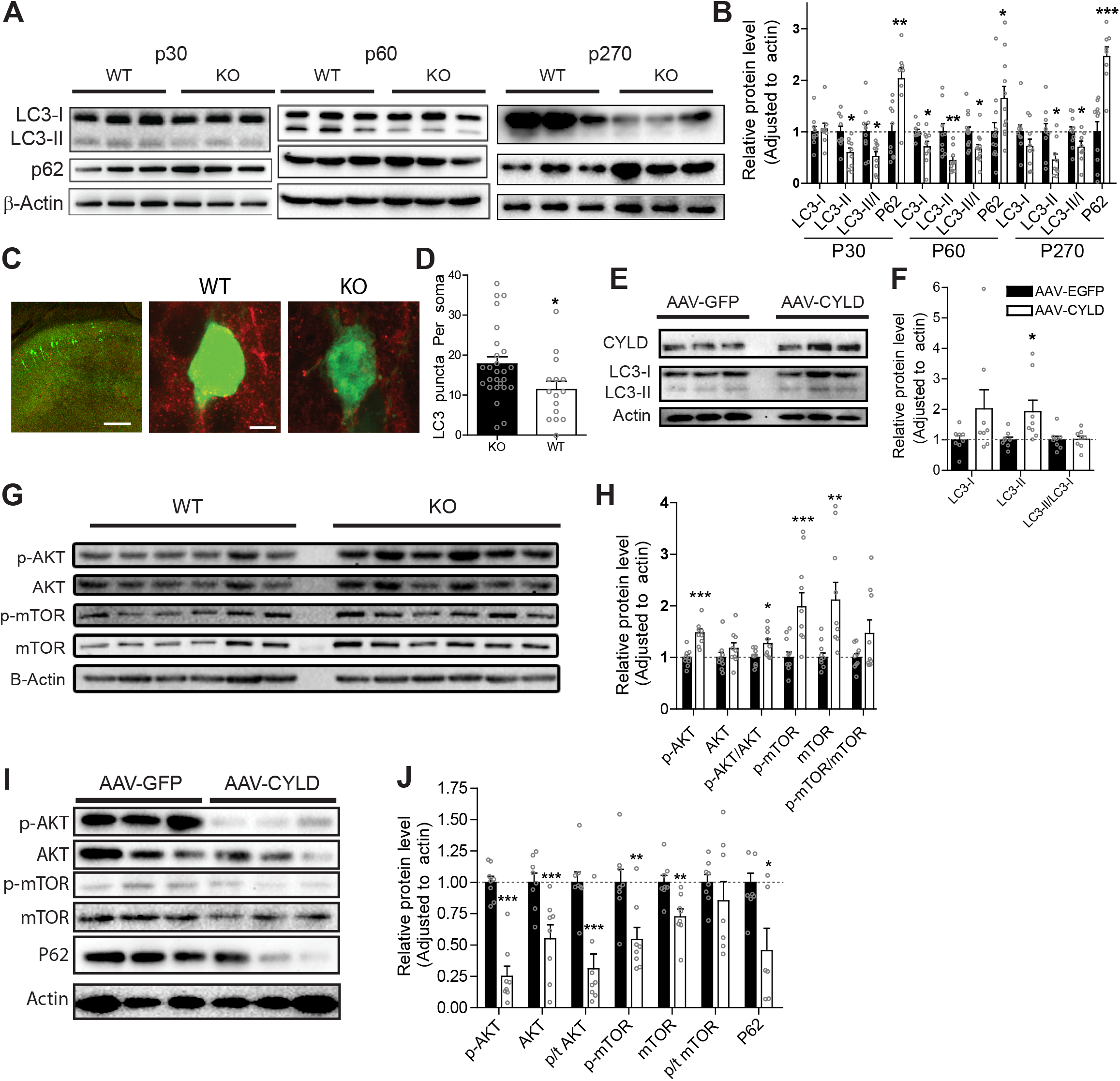
Impaired autophagy and heightened Akt-mTOR signaling in CYLD KO hippocampus. (A) Representative LC3 and p62 blots on hippocampal lysates prepared from p30, p60, and p270 WT and CYLD KO mice. (B) Quantifications from (A). Protein levels were first adjusted to β-actin and then normalized to respective WT levels. n = 10-12 mice/group. (C) Representative LC3 staining on hippocampal sections from CYLD WT and KO brains injected with AAV2-hSyn-EGFP and stained for LC3 (red). Scale bars: 200 μm (left) and 5 μm (right). (D) Quantifications of LC3 puncta from (C). LC3 puncta numbers per soma from WT and KO neurons infected with AAV-EGFP and stained for LC3 were quantified. n = 26 (WT) and 15 (KO). (E) Representative LC3 blots on lysates prepared (DIV21) from cultured hippocampal neurons infected (DIV7) with AAV-EGFP or AAV-CYLD. (F) Quantifications of (E). n = 8/group. (G) Representative total Akt, p-Akt (T308), total mTOR, p-mTOR (S2448), and β-actin blots on hippocampal lysates prepared from WT and CYLD KO mice. (H) Quantifications of (G). n = 9/group. (I) Representative blots on lysates prepared (DIV21) from cultured hippocampal neurons infected (DIV7) with AAV-EGFP or AAV-CYLD. (J) Quantifications of (I). n =6/group for p62; n = 8/group for all others. *P < 0.05, **P < 0.01, *** P < 0.001; Two-tailed unpaired t-tests.

A master inhibitor of autophagy is mTOR, the catalytic subunit of the mTORC1 complex, which upon activation via phosphorylation at S2448 inhibits the autophagy-initiating Ulk1 (Unc-51-like kinase) complex [48-50]. Phospho-mTOR level was significantly elevated in both hippocampal (Figure 5G and 5H) and cortical (Figure S4C and S4D) lysates from CYLD KO, suggesting hyperactive mTOR in mutant brains. Akt/PKB, an upstream activator of mTOR [49, 50], undergoes K63-linked polyubiquitination to promote its activation [51]. Akt is a direct substrate of CYLD which inhibits Akt by deubiquitination in non-neuron cells [9]. We found a significant increase of phosphorylated (T308), but not total levels of Akt in CYLD KO hippocampal (Figure 5G and 5H) and cortical (Figure S4C and S4D) lysates. Conversely, overexpressing CYLD in cultured hippocampal neurons produced a striking reduction of both total and phosphorylated levels of Akt and mTOR, accompanied with significantly reduced p62 (Figure 5I and 5J). These results reveal an Akt-mTOR-autophagy signaling axis that is potently regulated by CYLD.

### CYLD drives spine elimination in an autophagy-dependent manner

We next tested the hypothesis that CYLD drives spine pruning via autophagy in cultured neurons by treating with inhibitors that act at various steps along the autophagy pathway (Figure 6). Consistent with above data, CYLD overexpression significantly reduced total, mushroom, stubby, and thin spines compared to DsRed-expressing control neurons (Figure 6A-6D). Treating cells with the Akt activator SC79, but not the vehicle control DMSO, did not affect spine densities in DsRed cells but prevented spine density reductions in CYLD-overexpressing neurons (Figure 6A and 6B). Similar results were obtained with autophagy blockers Wortmannin and Bafilomycin A1, both blocking autophagosome-lysosome fusion, restoring total and mushroom spines to levels of DsRed/DMSO control (Figure 6A, 6C, and 6D). These data support that CYLD prunes synapses via an Akt- and autophagy activation-dependent mechanism.

**Figure 6.**
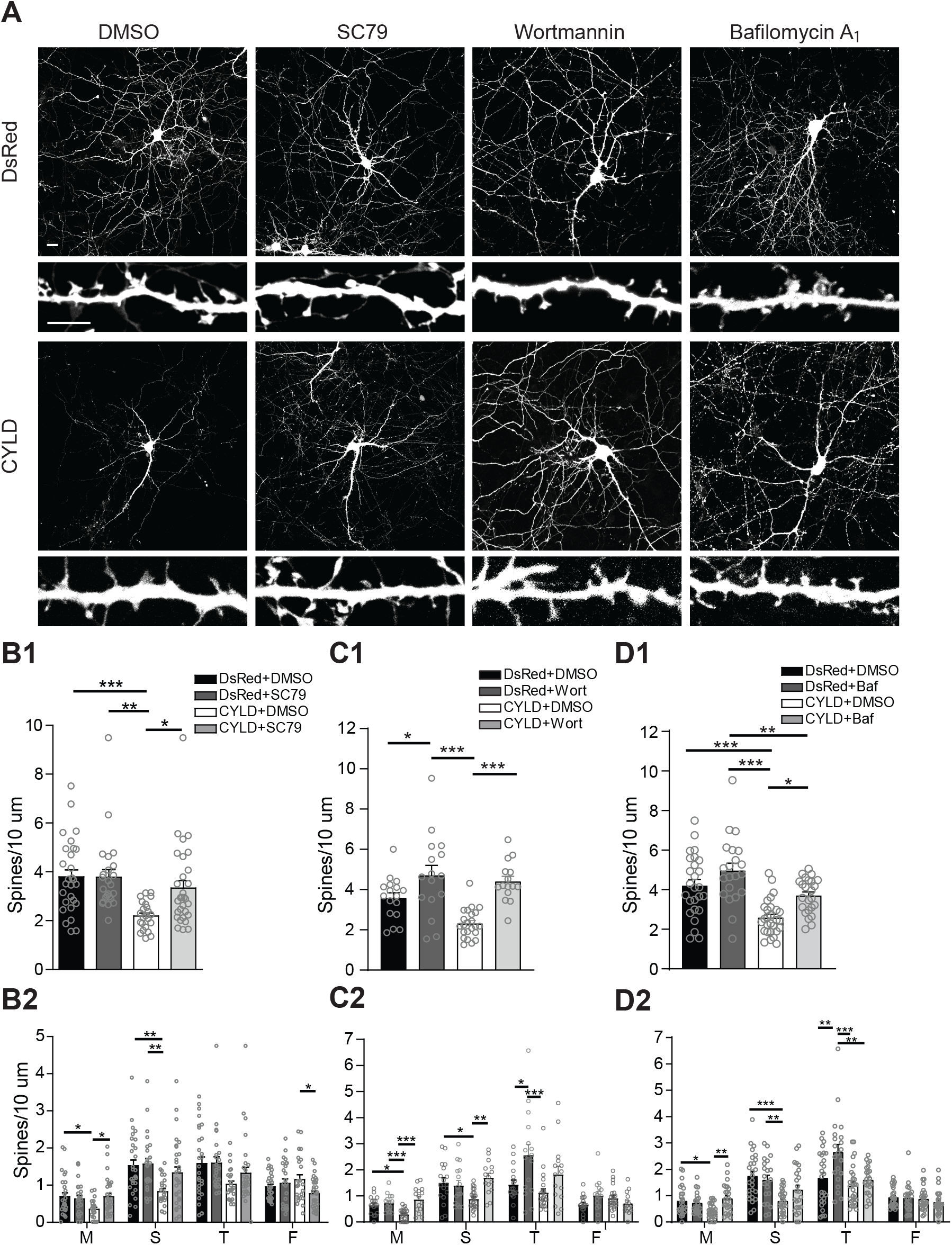
CYLD drives spine elimination via Akt and autophagy. (A) Representative whole-neuron (top) and primary dendrite (bottom) images from cultured hippocampal neurons transfected with DsRed or DsRed/CYLD, and treated with either DMSO (vehicle), SC79 (15 μM), Wortmannin (100 nM) or Bafilomycin A_1_ (10 nM) from DIV12-19. Scale bars: 20 and 5 μm. (B-D) Quantifications of SC79, Wortmannin (Wort), and Bafilomycin A1 (Baf) effects in spine densities. Total (B1, C1, and D1) and classified (B2, C2, and D2) spine densities are quantified from (A). n = 23-30 cells (SC79), 15-23 (Wort) and 21-28 (Baf) per group from 3 independent experiments. *P < 0.05, **P < 0.01, *** P < 0.001; One-way ANOVA with Bonferroni’s multiple comparison post-hoc test.

### Enhancing autophagy in CYLD KO rescues spine density, mEPSCs, and LTD

We finally examined if enhancing autophagy could restore the synaptic deficits in CYLD KO mice (Figure 7). Mice were intraperitoneally administered rapamycin or saline daily from p20-p28 and sacrificed for assessments on p30 (Figure 7A). For spine analysis, we also performed stereotaxic injections of AAV2-hSyn-EGFP to the CA1 hippocampus of WT and KO mice at P14/15 to label neurons (Figure 7B), which was then followed by the rapamycin regimen. As expected, there was increased autophagy flux (with increasing LC3-II) and decreased GluA1 level in rapamycin-treated KO mice compared to saline controls (Figure S5A.). Rapamycin treatments (1.5 mg/kg) did not affect WT spine densities compared to saline, but restored mushroom and stubby spine densities in KO to WT control levels (Figure 7C and 7D). Rapamycin at both low (1.5mg/kg/day) and medium (3mg/kg/day) doses also fully restored the enhanced mEPSC frequency in KO neurons to WT levels, where rapamycin did not have effects compared to saline (Figure 7E and 7F). Rapamycin treatments did not affect mEPSC amplitude compared to saline under any conditions (Figure 7E and 7F). Finally, rapamycin at 1.5/mg/kg also restored the lost LTD in KO mice to WT levels (Figure 7G and 7H). Together, these results provide strong evidence that reduced autophagy in KO mice underlies their impaired spine density, synaptic strength, and LTD deficits.

**Figure 7.**
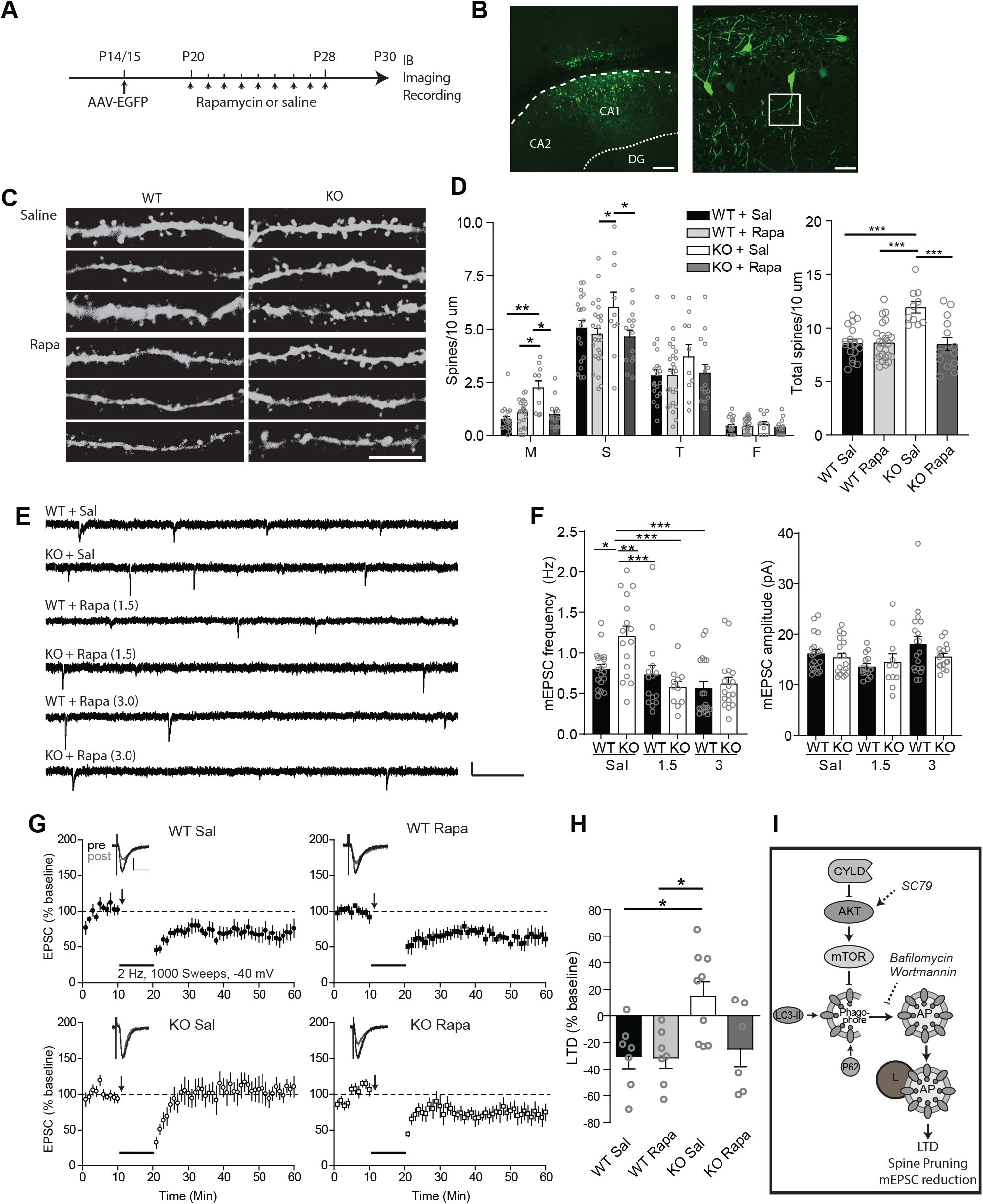
Rapamycin rescues spine density, synaptic efficacy, and LTD in CYLD KO mice. (A) Rapamycin treatment and analysis timeline. (B) Representative images showing AAV-EGFP infected neurons from a hippocampal slice. White box indicates the region of interest (first branch of the primary apical dendrite) for spine imaging. Scale bars: 200 μm (left) and 50 μm (right). (C) Representative dendritic segments from the first branch of CA1 hippocampal neurons prepared from CYLD WT and KO mice, treated with either saline or 1.5 mg/kg/day rapamycin. Scale bar: 5 μm. (D) Quantifications of spine densities from (C). Sal, saline; Rapa, Rapamycin. n = 11-19 cells per group from 4-6 mice per group. (E) Representative mEPSC traces from CYLD WT and KO mice treated with saline, 1.5 mg/kg/day rapamycin, or 3 mg/kg/day rapamycin. Scale bar: 10 pA, 500 ms. (F) mEPSCs quantifications from (E). n = 14-19 cells per group from 3-4 mice per group. (G) LTD rescue by rapamycin (1.5 mg/kg/day). Insets are representative EPSCs 5 min before (pre) and 40 min after (post) LTD induction. Scale bars: 50 pA, 25 ms. n= 7 (WT Sal), 7 (WT Rapa), 9 (KO Sal), and 6 (KO Rapa). (H) LTD quantification, calculated from 20-40 minutes post-induction. n= 6-7 cells/group. (I) Working model. CYLD facilitates induction of neuronal autophagy via inhibition of Akt-mTOR, driving spine pruning and LTD. Dashed lines indicate the effect of different drugs used. AP, autophagosome; L, lysosome. *P < 0.05, **P < 0.01, *** P < 0.001; One-way ANOVA with Bonferroni’s multiple comparison post-hoc test.

## DISCUSSION

In this study, we report that the K63-specific, FTD-linked DUB CYLD is a critical regulator of synaptic maintenance and plasticity by acting on the Akt-mTOR-autophagy axis (Figure 7I). We show that CYLD-deficient mice display aberrant locomotor, social, affective, and cognitive behaviors, accompanied with impaired synaptic numbers, efficacy, and plasticity. We further demonstrate that compromised neuronal autophagy resulting from heightened Akt-mTOR signaling underlies the synaptic function and plasticity deficits in mutant mice.

Based on its interactions with several autophagy receptors (i.e. p62/SQSTM1, Optineurin, and TBK1) [45, 46, 52] and HDAC6 [53], a protein that regulates autophagosome transport and maturation [54], CYLD is speculated to act as a “brake” on autophagy. Although mechanisms are unknown or unrelated, two recent studies support an inhibitory role for CYLD in autophagy in non-neuronal cells: CYLD has been shown to inhibit fusion of autophagosome with lysosome in HEK293 cells [18] and suppress autolysosome efflux in cardiomyocytes [55]. However, the role of CYLD in intact brains has not been investigated. Our study provides evidence that CYLD enhances autophagy flux in mouse hippocampus and cortex, mediated by suppression of Akt and mTOR activity. This might be unexpected but not entirely surprising due to diverse CYLD interactors or substrates that can act on different nodes of the autophagy cycle differentially, which can further vary depending on cell types, subcellular compartments, and cargo types. Regardless of mechanism details, our results indicate that in intact brain, a net effect of CYLD is to facilitate autophagy flux.

In neurons, the autophagosome is believed to primarily form at the distal axon then is retrogradely transported along microtubules to the lysosome-enriched soma for degradation, and roles of autophagy in presynaptic functions such as neurotransmitter release and cargo degradations, have been extensively investigated [29, 32, 56]. In contrast, postsynaptic autophagy is less well studied [30, 32]. However, recent studies reveal the presence and activity-dependent trafficking of lysosomes as well as synaptic activity-regulated autophagic vacuole motility in dendrites and spines [57, 58]. Furthermore, postsynaptic (presumably) autophagy promotes developmental pruning of dendritic spine in the cortex [37], contributes to NMDAR-dependent LTD and LTP [33-36] in the hippocampus, and mediates BDNF-regulated synaptic plasticity and memory in response to nutritional stress [59]. Nevertheless, much of the molecular mechanisms in postsynaptic compartments remain elusive [30, 32]. Given that most CYLD localizes at postsynaptic sites and postsynaptic mechanisms mediate the synaptic strength and plasticity deficits in CYLD KO mice, our study uncovers a novel signaling mechanism, CYLD-Akt-mTOR, that regulates postsynaptic autophagy.

Our results indicate that a physiological function of CYLD is to limit synapse strength and plasticity. CYLD promotes synapse elimination (but see [60]), likely by pruning/destabilizing existing spines from mature neurons in a DUB dependent manner rather than inhibiting synapse formation. Consistently, in the absence of CYLD, KO mice exhibit more abundant synapses, higher synaptic protein levels, and increased synaptic efficacy. CYLD-deficient synapses also show impaired NMDAR-dependent LTD (and to a lesser degree, LTP) with intact mGluR1-LTD (presumably because mTOR activity is not compromised), suggesting that CYLD plays an important role in shaping synaptic properties in addition to regulating synaptic density. Importantly, the spine morphology, mEPSC, and LTD deficits in CYLD mutant mice are rescued by restoring autophagy *in vivo*, supporting that decreased autophagy is behind these phenotypes. Overall, our results demonstrate that CYLD promotes synapse removal and mediates NMDAR-LTD via modulation of the Akt-mTOR signaling pathway through autophagy. However, it is worth noting that mTOR can also regulate synapse remodeling via other downstream cascades independent of autophagy, e.g. protein translation-dependent mechanisms [48, 61-63]. Furthermore, other CYLD substrates besides Akt, such as PSD-95 [15], β-catenin [64], and components in the NF-κB pathway [65-67] of which CYLD is a critical component [68-70], can regulate synapse remodeling and plasticity. Nevertheless, these potential downstream mechanisms and the Akt-mTOR-autophagy mechanism identified here are not mutually exclusive in regulating synapse remodeling and plasticity.

Our study establishes that CYLD is required to maintain an array of behaviors. The anxiety-like behavior observed in CYLD KO mice is consistent with a recent study [71]. In comparison, the role of CYLD in depression and social behavior has not been previously demonstrated. Furthermore, we show that CYLD is critical in cognitive flexibility, as CYLD KO mice displayed deficits in selecting an optimal strategy during reversal learning. This deficit can be explained at least partially by impaired LTD in mutant mice as hippocampal NMDAR-LTD is thought to mediate behavioral flexibility [72].

Recently, three rare variants of the *CYLD* gene, predicted to have high pathogenic potentials, were identified in FTD/ALS patients, placing *CYLD* as the newest member of the FTD/ALS-causing gene family [18, 20]. FTD is characterized by striking changes in personality, loss of empathy, apathy, disinhibition, and language disability at early-mid stages, and general cognitive deteriorations at later stages [73], likely due to synaptic and circuit dysfunctions. In addition, CYLD is a central regulator of immune signaling [69, 70], dysregulation of which is associated with many psychiatric and neurological diseases, which all involve synapse impairments and circuit mis-wiring [61, 74, 75]. Our findings thus provide a foundation for future investigation of synaptic and circuit deficits in neurological and psychiatric disease.

## MATERIALS AND METHODS

### Animals

CYLD-deficient mice were generated previously [11]. CYLD wildtype (WT), heterozygote, and homozygote (KO) mice were bred in-house by crossing CYLD heterozygous males and females, and genotyped using PCR. C57BL/6J mice were obtained from Jackson Laboratory (Bar Harbor, ME, USA) and bred in-house. Timed pregnant (E18) Wistar Rats were purchased from Charles River Laboratories (Wilmington, MA, USA). Mice were housed in same-sex groups of 2-5 under a 12h light/dark cycle with ad-libitum access to food and water. All animal procedures were approved by the SUNY Upstate or Icahn School of Medicine at Mount Sinai Institutional Animal Care and Use Committees.

### Plasmids

Wildtype CYLD, hCYLD, and CYLD C601A mutant constructs were generated using the pLL3.7 lentiviral vector backbone under the CMV promoter with an EGFP reporter [15]. Myc-hCYLD and Myc-CYLD C601A AAV plasmids were generated using the pAAV backbone under the hSyn promoter. pAAV-hSyn-DsRed was a gift from Edward Callaway (Addgene plasmid # 22907). Plasmids were transformed into XL10-GOLD ultracompetent E. coli (Agilent) according to the manufacturer’s protocol. Plasmid DNA was isolated using a Plasmid Midiprep Kit (Thermo Fisher) and stored in TE buffer (Thermo Fisher). Concentration and purity were recorded on a NanoDrop 1000 (Thermo Fisher). All constructs were confirmed by DNA sequencing and protein expression before use.

### Antibodies

Primary antibodies were used as follows. Western blot: Akt (1:1000; Cell Signaling, 9272), p-AKT [T308] (1:1000; Cell Signaling, 9275), CYLD (1:1000; Cell Signaling, 8462), mTOR (1:1000; Cell Signaling, 2983), p-mTOR [S2448] (1:1000; Cell Signaling, 5536), p62 (1:1000; Cell Signaling, 5114), LC3-II (1:1000; Cell Signaling, 38685), PSD95 (1:1000; Cell Signaling, 2507), GluA1 (1:1000, Cell Signaling, 13185), GluA2 (1:1000; Abcam, AB20673), GluN1 (1:1000; Cell Signaling, 5704), GluN2A (1:1000; Abcam, ab124913), GluN2B (1:1000; Abcam, ab254356), β-Actin (1:2000; Santa Cruz, SC-47778), Goat anti-Mouse IgG (H+L) HRP (1:2000; Invitrogen, A16066), Goat anti-Rabbit IgG (H+L) HRP (1:2000; Invitrogen, 32460). Staining: LC3-II (1:200; Santa Cruz, SC-376404), Alexa Fluor 647 Goat anti-Rabbit IgG (1:500; Thermo Fisher, A-21244).

### Cell Culture, Transfections, and Treatments

Primary hippocampal neurons were isolated from either E18 rat embryos or P0 mouse pups using a Papain Dissociation Kit according to the manufacturer’s protocol (Worthington Biochemistry). Cells were plated on poly-D-lysine (Sigma Aldrich) coated coverslips in 24-well dishes at medium density (∼60,000 cells/coverslip). Cells were cultured in Neurobasal medium (Thermo Fisher), and supplemented with 1X GlutaMax (Thermo Fisher), 1X B27 (Gibco), and 200 U/mL Penn/Strep (Thermo Fisher), and incubated at 37°C in 5% CO_2_. Medium was changed every 3-4 days with 50% fresh medium. Cells were cotransfected with hSyn-DsRed and the indicated construct on DIV 7 or DIV 19 (for older neurons) with 1 μg total purified plasmid DNA via CalPhos Mammalian Transfection Kit (Takara Bio) according to the manufacturer’s protocol.

Successful transfection was confirmed 24 hours later by fluorescence microscopy. For drug treatments, cells were treated with either SC79 (15 μM; Sigma), wortmannin (100 nM; Sigma), bafilomycin (10 nM; Sigma), or an equal volume of DMSO as vehicle control. Media was replaced daily with 50% untreated conditioned media collected previously from the same batch of neurons, and 50% fresh media containing each drug. Drug-treated samples were processed in parallel. DMSO controls that were processed simultaneously with multiple drug treatments were included in the data for each applicable group.

### In Vitro Virus Preparation and Infection

AAV particles used for in vitro infection were packaged by Vigene Biosciences. To infect cultured neurons, concentrated virus was added to DIV7 neurons at an MOI of ∼5,000 overnight. The next day, media was replaced with 50% fresh, 50% conditioned neurobasal media and cells were grown to DIV21 for analysis.

### Stereotaxic Surgery and In Vivo Labelling

For in vivo dendritic spine analysis, P14-15 WT and CYLD KO mice were anesthetized and injected with 0.2 μL AAV2-hSyn-EGFP (Addgene #50465-AAV2) into the CA1 hippocampus (AP: -1.8 mm, ML: +/-1.5 mm, DV: -1.3 mm; 0.05 μm/min) with a stereotaxic apparatus (Stoelting). Mice were injected intraperitoneally with 1.5 mg/kg or 3.0 mg/kg rapamycin (Sigma Aldrich) or saline each day from P20-P28. On P30, mice were anesthetized with ketamine/xylazine cocktail for cardiac perfusions. Mice were perfused with Phosphate Buffered Saline (PBS) followed by 4% paraformaldehyde (PFA) in PBS. Brains were post-fixed overnight at 4°C in 4% PFA with 5% sucrose. Brains were then embedded in low-melting temperature agarose and sliced transversely at 100 μm on a Vibratome 1000 (TPI). Slices were stored in 4% PFA in 5% sucrose until imaging. For LC3-II staining in vivo, fixed slices were blocked in blocking buffer (1X PBS with 5% normal donkey serum (Sigma) and 0.3% Triton X-100 (Sigma)) for 1 hour, incubated with primary antibody in blocking buffer overnight, rinsed 3X with PBS, and incubated with secondary antibody in dilution buffer (1X PBS, 1% BSA, 0.3% Triton X-100) for 1 hour.

### Immunoblotting

Cortical and hippocampal tissues were rapidly (under one minute) extracted on ice, snap frozen on dry ice, and stored at -80°C until lysis. Tissue was homogenized in ice-cold radioimmunoprecipitation assay (RIPA) buffer (50 mM Tris-HCL (pH 7.4), 1% NP-40, 0.1% SDS, 0.25% deoxycholic acid, 150 mM NaCl, 1 mM EDTA) supplemented with 1 mM PFSF, 1X protease inhibitor cocktail (Roche), and 1X Phosphatase Inhibitor Cocktail 3 (Sigma Aldrich). Cultured neurons were rinsed with ice-cold PBS and removed from coverslips in ice-cold RIPA buffer with a cell scraper. Supernatants were collected by centrifugation for western blotting. Lysates were mixed with SDS loading buffer and heated to 95 °C for 5 minutes before loading on a 7.5% or 15% (for resolving LC3-II) SDS-PAGE gel. Proteins were transferred to a PVDF membrane (Bio-Rad) and blocked in 3% BSA in Tris Buffered Saline with 0.1% Tween 20 (TBS-T; Santa Cruz). PVDF membranes were incubated with primary antibodies at 4°C overnight, and secondary antibodies at room temperature for 1 hour. For resolution of multiple proteins, blots were stripped between probes with Membrane Stripping Buffer-3 (Boston BioProducts). Signals were detected using SuperSignal West Femto Maximum Sensitivity Substrate Kit (Pierce) on a ChemiDoc MP (BioRad). Relative protein band intensity was quantified within linear range using ImageJ (NIH).

### Confocal Microscopy and Image Analysis

Whole-cell and dendritic images were acquired using a Leica TSC SP5 Spectral Confocal Microscope (Leica Microsystems) with a 40x or 63x (with 4x digital zoom) objectives at a resolution of 1024 × 1024 pixels. Brain slices and coverslips were mounted in ProB Antifade Mounting Reagent (Thermo Fisher) and imaged and analyzed blinded to sample, genotype, and condition. For brain slices, CA1 hippocampal pyramidal neurons were visually identified and imaged. For cultured cells, pyramidal neurons were randomly selected for imaging. Images were taken along the Z-axis at a total thickness of 1 μm (for dendrite segments), 3 μm (for whole-cell images of cultured neurons), or 5 μm (for whole-cell images of neurons in brain slices). Dendrites were imaged at the first branch of the primary apical dendrite, and every spine along the >25 μm length was measured using ImageJ. Spine classification was automatically sorted in Excel (Microsoft) by the following criteria: mushroom (head width:neck length >1.5), stubby (total length < 1 μm, head width:neck length < 1.5), thin (neck length >1 μm, head width:neck length < 1.5), filopodia (no discernible head) [76].

### Electrophysiology

For hippocampal slice recording, male or female WT and CYLD KO mice (P20-P30) were sacrificed and brains rapidly removed and sectioned transversely at 300 or 350 μm with a S1200 vibratome (Leica) in ice-cold, oxygenated (95% O_2_, 5% CO_2_; Airgas) sucrose cutting solution containing 75 mM sucrose, 87 mM NaCl, 2.5 mM KCl, 1.25 NaH_2_PO_4_, 7 mM MgCl_2_, 25 mM NaHCO_3_, 10 mM glucose, 0.5 mM CaCl_2_, and 1.3 mM ascorbic acid. Slices were placed in a preincubation chamber at 37°C for 45 minutes in artificial cerebrospinal fluid (aCSF) containing 126 mM NaCl, 18 mM NaHCO_3_, 2.5 mM KCl, 2.4 mM MgCl_2_, 1.2 mM CaCl_2_, 1.2 mM NaH_2_PO_4_, and 11 mM D-glucose [77]. Slices were then transferred to room temperature for the remainder of the experiment. Following incubation, slices were placed in a recording chamber containing oxygenated ACSF with 100 μM picrotoxin at 32°C. Medial prefrontal cortex (mPFC, containing anterior cingulate cortex and prelimbic cortex) slices were prepared from P30 or P60 mice as described in our previous publications [78-81]. Briefly, 300 μm coronal slices were cut in ice-cold oxygenated aCSF containing 126 mM NaCl, 25 mM NaHCO_3_, 2.5 mM KCl, 2.5 mM CaCl_2_, 1.2 mM MgCl_2_, 1.2 mM NaH_2_PO_4_, and 11 mM D-glucose, and following incubation, transferred to a recording chamber perfused with aCSF (saturated with 95% O_2_ and 5% CO_2_) for mPFC recording.

Whole-cell patch-clamp recordings were performed on visually identified CA1 pyramidal neurons in the hippocampus or layer V pyramidal neurons in the mPFC. For CA1 current-clamp recordings, patch pipettes (3.5-5.5 MΩ) were filled with internal solution containing 128 K-gluconate, 8 mM NaCl, 0.4 mM EGTA, 2 mM Mg-ATP, 5 mM QX-314, 0.3 mM Na-GTP, 10 mM HEPES, adjusted to pH 7.3. For mPFC and CA1 voltage-clamp recording, pipettes (3.5-5.5 MΩ) were filled with internal solution containing 142 mM Cs-gluconate, 8 mM NaCl, 10 mM HEPES, 0,4 mM EGTA, 2.5 mM QX-314, 2 mM Mg-ATP, and 0.25 mM GTP-Tris, adjusted to pH 7.2. All recordings were made at 32°C with a temperature controller (Warner Instruments). Drugs were delivered to the extracellular bath with a perfusion system (Harvard Apparatus).

For miniature excitatory postsynaptic current (mEPSC) recordings, 1 μM TTX was included in the bath to block voltage-gated sodium channels. Neurons were voltage-clamped at -60 mV for 10-min recordings. For recordings of evoked excitatory postsynaptic currents (eEPSCs) or excitatory postsynaptic potentials (eEPSPs), Schaffer collateral afferents in the stratum radiatum were stimulated using a concentric bipolar electrode (FHC) to evoke a response in CA1 hippocampal neurons. Only cells displaying a single, monosynaptic response were included in analysis. For paired-pulse ratio (PPR), a series of two consecutive pulses were delivered at desired inter-pulse intervals (20 ms, 50 ms, 100 ms and 200 ms). For presynaptic release probability assessment using MK-801, cells were held at -30 mV with CNQX (20 μM) added in bath to block AMPAR-EPSCs. Stimuli were delivered at 0.033 Hz. After 10 min recording of a stable baseline NMDAR-EPSCs, MK-801 (20 μM) was applied. Input-output (I/O) curve was determined on neurons voltage-clamped at -60 mV and stimulated with a series of stimuli starting at 80 pA and increasing by 80 pA per step, or 100 pA per step above 1000 pA. For synaptic summation analysis, cells were maintained at their resting membrane potential in current clamp mode. A train of 15 pulses at 20 Hz was delivered to trigger a train of eEPSPs followed by a test pulse. The stimulation intensity was controlled so that the amplitude of the first eEPSP was 2 to 6 mV.

For synaptic plasticity experiments, baseline eEPSC was established by holding cells at -60 mV, and presynaptic stimuli (200 μs duration at 0.033 Hz) were delivered with stimulation intensity controlled such that the amplitude of the baseline eEPSC was 150-250 pA. After obtaining stable eEPSCs for 10 min, LTP was induced by pairing presynaptic stimulation of 120 pulses at 2 Hz with depolarization of postsynaptic cells at +30 mV. LTD was induced by pairing 1000 presynaptic stimuli (at 2 Hz) with postsynaptic depolarization at -40 mV. mGluR1 was induced by bath-applied DHPG (75 μM) for 10 min. Plasticity was measured as the eEPSC amplitude after induction, divided by average baseline amplitude. Series resistance was monitored throughout whole-cell recordings, and data were discarded if the resistance changed by >15%.

Signals were collected with a MultiClamp 700B amplifier (Molecular Devices) and digitized at 10 kHz on a Digidata 1440A (Molecular Devices), filtered at 1 kHz. mEPSCs were analyzed using MiniAnalysis software (Synaptosoft), and eEPSC/eEPSPs were analyzed by pClamp 10 software (Molecular Devices).

### Behavioral Tests

Age-matched adult male and female mice (3-4 months old) were acclimated to the behavior rooms for at least 30 minutes in their home cage prior to all behavior testing. All testing chambers were cleaned between trials and subjects with REScue veterinary cleaner (Rescue Disinfectants). All behavioral assays were carried out and scored blind to genotype.

#### Open-field test

Mice were placed in a 40 cm x 40 cm x 30 cm open-field chamber (Med Associates) for one hour. Total distance and time in the peripheral and center were assessed by AnyMaze (Stoelting).

#### Light-dark box test

Mice were placed into the dark box insert in an open field chamber (Med Associates). Mice were allowed to move freely in the entire apparatus for 10 minutes. Time spent in each chamber, latency to enter the light portion of the apparatus, and the number of crossings between chambers was recorded.

#### Elevated plus maze test

Mice were placed on an elevated plus maze (Med Associates) containing two ‘open’ arms with no walls, and two ‘closed’ arms with 30.5 cm tall walls surrounding the arms. Mice were allowed to explore the apparatus for 5 minutes. Time in each arm and entries to each arm were recorded by an overhead camera and scored automatically with AnyMaze.

#### Forced-swim test

Mice were placed in a 50 cm tall x 18 cm diameter glass beaker containing fresh 22°C tap water filled to 24 cm to ensure mice could not climb out or touch the bottom of the beaker. Mice were allowed to swim freely for an induction phase of 5 minutes. 24 hours later, mice were placed back in the beaker for 5 minutes and locomotion was recorded with AnyMaze. Immobility was scored as a lack of climbing, diving, or swimming beyond what was required to stay above water for 2 or more seconds.

#### Tail suspension test

Mice were suspended by the tail using tape on a suspension bar for 5 minutes while recorded with a camera. Latency to freeze and time spent immobile were evaluated blindly by an observer. Animals showing tail climbing behaviors were removed from further analysis.

#### Three-chamber sociability test

Mice were placed into the center of a modular chamber (Med Associates) 120 cm x 40 cm x 40 cm, with one modular chamber on each side, separated by a plexiglass divider. Mice were acclimated to the center chamber for 5 minutes before being allowed to explore the remaining segments for another 10 minutes. The left and right chambers each contained a wire cup which was either empty or contained a novel age-and sex-matched non-littermate mouse. Time spent interacting with the empty cup or stranger mouse was manually scored. Time in each chamber was automatically scored by AnyMaze.

#### Y-maze

To test working memory, mice were placed in a Y maze apparatus (Med Associates) and allowed to explore for 5 minutes. Alternations were scored automatically by AnyMaze as sequential explorations of all three arms, without returning to a previously explored arm. Alternation percentage was defined as the number of complete alternations divided by the number of total trials multiplied by 100.

#### Barnes maze

To assess learning and memory, mice were briefly trained to locate the escape tunnel of a circular Barnes Maze table (48” diameter, 20 holes evenly spaced along the edge, Med Associates). The escape tunnel was large enough to accommodate the mouse. The next day, mice were allowed to explore the Barnes Maze for 3 minutes, with their time spent until entering the escape tunnel and number of non-escape tunnel explorations recorded with AnyMaze. The test was ended either at the end of three minutes, or when both of the animal’s forelimbs had entered the escape tunnel. Animals that did not locate the escape tunnel by the end of the test were guided to the escape tunnel. This was repeated three times per day for 4 days. To assess behavioral flexibility, the escape tunnel was relocated to the hole 180° from the original position, on the opposite side of the table. Mice were allowed 5 minutes per day, once per day to locate the new location of the escape tunnel.

### Statistical Analysis

All data is represented as mean ± SEM. Comparisons involving two groups were analyzed with a two-tailed unpaired Student’s t test. For data containing more than two groups, a one-factor ANOVA was performed followed by appropriate multiple comparison post-hoc test, with a significance threshold set at p = 0.05. For cumulative probability tests, the Komogorov-Smirnov test was used. All statistical analyses were performed using Prism 7 (GraphPad).

## ACKNOWLEDGEMENTS

We thank members of the laboratory for critiques and comments. This work was supported by NIH Grants MH106489, NS093097, RR026761 (to W.-D.Y.) and RR000168 (to the New England Primate Research Center/Harvard Medical School).

## CONFLICT OF INTEREST

The authors declare no conflict of interest.

## Notes

### Competing Interest Statement

The authors have declared no competing interest.

## REFERENCES

1. Komander, D., et al., The structure of the CYLD USP domain explains its specificity for Lys63-linked polyubiquitin and reveals a B box module. Mol Cell, 2008. 29(4): p. 451–64.

2. Bignell, G.R., et al., Identification of the familial cylindromatosis tumour-suppressor gene. Nat Genet, 2000. 25(2): p. 160–5.

3. Mukhopadhyay, D. and H. Riezman, Proteasome-independent functions of ubiquitin in endocytosis and signaling. Science, 2007. 315(5809): p. 201–5.

4. Chen, Z.J. and L.J. Sun, Nonproteolytic functions of ubiquitin in cell signaling. Mol Cell, 2009. 33(3): p. 275–86.

5. Husnjak, K. and I. Dikic, Ubiquitin-binding proteins: decoders of ubiquitin-mediated cellular functions. Annu Rev Biochem, 2012. 81: p. 291–322.

6. Brummelkamp, T.R., et al., Loss of the cylindromatosis tumour suppressor inhibits apoptosis by activating NF-kappaB. Nature, 2003. 424(6950): p. 797–801.

7. Kovalenko, A., et al., The tumour suppressor CYLD negatively regulates NF-kappaB signalling by deubiquitination. Nature, 2003. 424(6950): p. 801–5.

8. Trompouki, E., et al., CYLD is a deubiquitinating enzyme that negatively regulates NF-kappaB activation by TNFR family members. Nature, 2003. 424(6950): p. 793–6.

9. Lim, J.H., et al., CYLD negatively regulates transforming growth factor-beta-signalling via deubiquitinating Akt. Nat Commun, 2012. 3: p. 771.

10. Massoumi, R., et al., Cyld inhibits tumor cell proliferation by blocking Bcl-3-dependent NF-kappaB signaling. Cell, 2006. 125(4): p. 665–77.

11. Reiley, W.W., et al., Regulation of T cell development by the deubiquitinating enzyme CYLD. Nat Immunol, 2006. 7(4): p. 411–7.

12. Zhang, J., et al., Impaired regulation of NF-kappaB and increased susceptibility to colitis-associated tumorigenesis in CYLD-deficient mice. J Clin Invest, 2006. 116(11): p. 3042–9.

13. Zajicek, A. and W.D. Yao, Remodeling without destruction: non-proteolytic ubiquitin chains in neural function and brain disorders. Mol Psychiatry, 2021. 26(1): p. 247–264.

14. Dosemeci, A., et al., CYLD, a deubiquitinase specific for lysine63-linked polyubiquitins, accumulates at the postsynaptic density in an activity-dependent manner. Biochem Biophys Res Commun, 2013. 430(1): p. 245–9.

15. Ma, Q., et al., Proteasome-independent polyubiquitin linkage regulates synapse scaffolding, efficacy, and plasticity. Proc Natl Acad Sci U S A, 2017. 114(41): p. E8760–E8769.

16. Thein, S., et al., CaMKII mediates recruitment and activation of the deubiquitinase CYLD at the postsynaptic density. PLoS One, 2014. 9(3): p. e91312.

17. Jin, C., et al., Shank3 regulates striatal synaptic abundance of Cyld, a deubiquitinase specific for Lys63-linked polyubiquitin chains. J Neurochem, 2019. 150(6): p. 776–786.

18. Dobson-Stone, C., et al., CYLD is a causative gene for frontotemporal dementia - amyotrophic lateral sclerosis. Brain, 2020. 143(3): p. 783–799.

19. Zhang, J., et al., Altered striatal rhythmic activity in cylindromatosis knock-out mice due to enhanced GABAergic inhibition. Neuropharmacology, 2016. 110(Pt A): p. 260–267.

20. Tabuas-Pereira, M., et al., CYLD variants in frontotemporal dementia associated with severe memory impairment in a Portuguese cohort. Brain, 2020.

21. Mizushima, N. and M. Komatsu, Autophagy: renovation of cells and tissues. Cell, 2011. 147(4): p. 728–41.

22. Klionsky, D.J., Autophagy: from phenomenology to molecular understanding in less than a decade. Nat Rev Mol Cell Biol, 2007. 8(11): p. 931–7.

23. Hara, T., et al., Suppression of basal autophagy in neural cells causes neurodegenerative disease in mice. Nature, 2006. 441(7095): p. 885–9.

24. Komatsu, M., et al., Loss of autophagy in the central nervous system causes neurodegeneration in mice. Nature, 2006. 441(7095): p. 880–4.

25. Nixon, R.A., The role of autophagy in neurodegenerative disease. Nat Med, 2013. 19(8): p. 983–97.

26. Yamamoto, A. and Z. Yue, Autophagy and its normal and pathogenic states in the brain. Annu Rev Neurosci, 2014. 37: p. 55–78.

27. Menzies, F.M., et al., Autophagy and Neurodegeneration: Pathogenic Mechanisms and Therapeutic Opportunities. Neuron, 2017. 93(5): p. 1015–1034.

28. Gao, F.B., S. Almeida, and R. Lopez-Gonzalez, Dysregulated molecular pathways in amyotrophic lateral sclerosis-frontotemporal dementia spectrum disorder. EMBO J, 2017. 36(20): p. 2931–2950.

29. Stavoe, A.K.H. and E.L.F. Holzbaur, Autophagy in Neurons. Annu Rev Cell Dev Biol, 2019. 35: p. 477–500.

30. Nikoletopoulou, V. and N. Tavernarakis, Regulation and Roles of Autophagy at Synapses. Trends Cell Biol, 2018. 28(8): p. 646–661.

31. Tomoda, T., K. Yang, and A. Sawa, Neuronal Autophagy in Synaptic Functions and Psychiatric Disorders. Biol Psychiatry, 2020. 87(9): p. 787–796.

32. Lieberman, O.J. and D. Sulzer, The Synaptic Autophagy Cycle. J Mol Biol, 2020. 432(8): p. 2589–2604.

33. Shehata, M., et al., Neuronal stimulation induces autophagy in hippocampal neurons that is involved in AMPA receptor degradation after chemical long-term depression. J Neurosci, 2012. 32(30): p. 10413–22.

34. Compans, B., et al., NMDAR-dependent long-term depression is associated with increased short term plasticity through autophagy mediated loss of PSD-95. Nat Commun, 2021. 12(1): p. 2849.

35. Shen, H., et al., Autophagy controls the induction and developmental decline of NMDAR-LTD through endocytic recycling. Nat Commun, 2020. 11(1): p. 2979.

36. Glatigny, M., et al., Autophagy Is Required for Memory Formation and Reverses Age-Related Memory Decline. Curr Biol, 2019. 29(3): p. 435–448 e8.

37. Tang, G., et al., Loss of mTOR-dependent macroautophagy causes autistic-like synaptic pruning deficits. Neuron, 2014. 83(5): p. 1131–43.

38. Levine, B. and G. Kroemer, SnapShot: Macroautophagy. Cell, 2008. 132(1): p. 162 e1–162 e3.

39. Saxton, R.A. and D.M. Sabatini, mTOR Signaling in Growth, Metabolism, and Disease. Cell, 2017. 169(2): p. 361–371.

40. Yan, J., et al., Activation of autophagy rescues synaptic and cognitive deficits in fragile X mice. Proc Natl Acad Sci U S A, 2018. 115(41): p. E9707–E9716.

41. Zucker, R.S. and W.G. Regehr, Short-term synaptic plasticity. Annu Rev Physiol, 2002. 64: p. 355–405.

42. Jakovcevski, M., et al., Neuronal Kmt2a/Mll1 histone methyltransferase is essential for prefrontal synaptic plasticity and working memory. J Neurosci, 2015. 35(13): p. 5097–108.

43. Kasai, H., et al., Structure-stability-function relationships of dendritic spines. Trends Neurosci, 2003. 26(7): p. 360–8.

44. Shen, W. and B. Ganetzky, Autophagy promotes synapse development in Drosophila. J Cell Biol, 2009. 187(1): p. 71–9.

45. Friedman, C.S., et al., The tumour suppressor CYLD is a negative regulator of RIG-I-mediated antiviral response. EMBO Rep, 2008. 9(9): p. 930–6.

46. Nagabhushana, A., M. Bansal, and G. Swarup, Optineurin is required for CYLD-dependent inhibition of TNFalpha-induced NF-kappaB activation. PLoS One, 2011. 6(3): p. e17477.

47. Wooten, M.W., et al., Essential role of sequestosome 1/p62 in regulating accumulation of Lys63-ubiquitinated proteins. J Biol Chem, 2008. 283(11): p. 6783–9.

48. Huber, K.M., et al., Dysregulation of Mammalian Target of Rapamycin Signaling in Mouse Models of Autism. J Neurosci, 2015. 35(41): p. 13836–42.

49. Hay, N. and N. Sonenberg, Upstream and downstream of mTOR. Genes Dev, 2004. 18(16): p. 1926–45.

50. Laplante, M. and D.M. Sabatini, mTOR signaling in growth control and disease. Cell, 2012. 149(2): p. 274–93.

51. Yang, W.L., et al., The E3 ligase TRAF6 regulates Akt ubiquitination and activation. Science, 2009. 325(5944): p. 1134–8.

52. Jin, W., et al., Deubiquitinating enzyme CYLD negatively regulates RANK signaling and osteoclastogenesis in mice. J Clin Invest, 2008. 118(5): p. 1858–66.

53. Wickstrom, S.A., et al., CYLD negatively regulates cell-cycle progression by inactivating HDAC6 and increasing the levels of acetylated tubulin. EMBO J, 2010. 29(1): p. 131–44.

54. Chin, L.S., J.A. Olzmann, and L. Li, Parkin-mediated ubiquitin signalling in aggresome formation and autophagy. Biochem Soc Trans, 2010. 38(Pt 1): p. 144–9.

55. Qi, L., et al., CYLD exaggerates pressure overload-induced cardiomyopathy via suppressing autolysosome efflux in cardiomyocytes. J Mol Cell Cardiol, 2020. 145: p. 59–73.

56. Luningschror, P. and M. Sendtner, Autophagy in the presynaptic compartment. Curr Opin Neurobiol, 2018. 51: p. 80–85.

57. Goo, M.S., et al., Activity-dependent trafficking of lysosomes in dendrites and dendritic spines. J Cell Biol, 2017. 216(8): p. 2499–2513.

58. Kulkarni, V.V., et al., Synaptic activity controls autophagic vacuole motility and function in dendrites. J Cell Biol, 2021. 220(6).

59. Nikoletopoulou, V., et al., Modulation of Autophagy by BDNF Underlies Synaptic Plasticity. Cell Metab, 2017. 26(1): p. 230–242 e5.

60. Li, J., et al., Tumor suppressor protein CYLD regulates morphogenesis of dendrites and spines. Eur J Neurosci, 2019. 50(4): p. 2722–2739.

61. Estes, M.L. and A.K. McAllister, Immune mediators in the brain and peripheral tissues in autism spectrum disorder. Nat Rev Neurosci, 2015. 16(8): p. 469–86.

62. Borrie, S.C., et al., Cognitive Dysfunctions in Intellectual Disabilities: The Contributions of the Ras-MAPK and PI3K-AKT-mTOR Pathways. Annu Rev Genomics Hum Genet, 2017. 18: p. 115–142.

63. Richter, J.D., G.J. Bassell, and E. Klann, Dysregulation and restoration of translational homeostasis in fragile X syndrome. Nat Rev Neurosci, 2015. 16(10): p. 595–605.

64. Tauriello, D.V., et al., Loss of the tumor suppressor CYLD enhances Wnt/beta-catenin signaling through K63-linked ubiquitination of Dvl. Mol Cell, 2010. 37(5): p. 607–19.

65. Kaltschmidt, B. and C. Kaltschmidt, NF-kappaB in the nervous system. Cold Spring Harb Perspect Biol, 2009. 1(3): p. a001271.

66. Meffert, M.K. and D. Baltimore, Physiological functions for brain NF-kappaB. Trends Neurosci, 2005. 28(1): p. 37–43.

67. Mei, S., et al., The ubiquitin-editing enzyme A20 regulates synapse remodeling and efficacy. Brain Res, 2020. 1727: p. 146569.

68. Chen, J. and Z.J. Chen, Regulation of NF-kappaB by ubiquitination. Curr Opin Immunol, 2013. 25(1): p. 4–12.

69. Komander, D., M.J. Clague, and S. Urbe, Breaking the chains: structure and function of the deubiquitinases. Nat Rev Mol Cell Biol, 2009. 10(8): p. 550–63.

70. Sun, S.C., Deubiquitylation and regulation of the immune response. Nat Rev Immunol, 2008. 8(7): p. 501–11.

71. Han, Y.Y., et al., Microglial activation in the dorsal striatum participates in anxiety-like behavior in Cyld knockout mice. Brain Behav Immun, 2020.

72. Collingridge, G.L., et al., Long-term depression in the CNS. Nat Rev Neurosci, 2010. 11(7): p. 459–73.

73. Neary, D., et al., Frontotemporal lobar degeneration: a consensus on clinical diagnostic criteria. Neurology, 1998. 51(6): p. 1546–54.

74. Boulanger, L.M. and C.J. Shatz, Immune signalling in neural development, synaptic plasticity and disease. Nat Rev Neurosci, 2004. 5(7): p. 521–31.

75. Hammond, T.R., S.E. Marsh, and B. Stevens, Immune Signaling in Neurodegeneration. Immunity, 2019. 50(4): p. 955–974.

76. Harris, K.M., F.E. Jensen, and B. Tsao, Three-dimensional structure of dendritic spines and synapses in rat hippocampus (CA1) at postnatal day 15 and adult ages: implications for the maturation of synaptic physiology and long-term potentiation. J Neurosci, 1992. 12(7): p. 2685–705.

77. Chaouloff, F., A. Hemar, and O. Manzoni, Acute stress facilitates hippocampal CA1 metabotropic glutamate receptor-dependent long-term depression. J Neurosci, 2007. 27(27): p. 7130–5.

78. Xu, T.X., et al., Hyperdopaminergic tone erodes prefrontal long-term potential via a D2 receptor-operated protein phosphatase gate. J Neurosci, 2009. 29(45): p. 14086–99.

79. Xu, T.X., et al., Amphetamine modulation of long-term potentiation in the prefrontal cortex: dose dependency, monoaminergic contributions, and paradoxical rescue in hyperdopaminergic mutant. J Neurochem, 2010. 115(6): p. 1643–54.

80. Ruan, H. and W.D. Yao, Cocaine Promotes Coincidence Detection and Lowers Induction Threshold during Hebbian Associative Synaptic Potentiation in Prefrontal Cortex. J Neurosci, 2017. 37(4): p. 986–997.

81. Ruan, H. and W.D. Yao, Loss of mGluR1-LTD following cocaine exposure accumulates Ca(2+)-permeable AMPA receptors and facilitates synaptic potentiation in the prefrontal cortex. J Neurogenet, 2021: p. 1–12.

