## Supplemental figures and legends for "Cylindromatosis Drives Synapse Pruning and Weakening by Promoting Macroautophagy through Akt-mTOR Signaling"

**Supplementary Figure Legends**

**Figure S1.** Unaltered working memory in CYLD KO mice. Fraction of alternation during a 5-minute Y Maze test of WT and CYLD KO mice. Alternation percentage is calculated as the number of complete alternations/total number of trials multiplied by 100.

**Figure S2. mEPSC analysis in mPFC.** (A) Representative mEPSC traces from layer V pyramidal neurons in the mPFC from P30 and P60 WT and CYLD KO mice. (B-C) Quantifications of mEPSC frequency (B) and amplitude (C). n = 30 (P30 WT), 19 (P30 KO), 18 (P60 WT), and 17 (P60 KO). Scale bar: 20 pA, 200 ms. * $p < 0.05$; two-tailed unpaired t-test.

**Figure S3. Synaptic protein blots in AAV-infected cultured neurons.** Representative western blots (A) of GluA1, GluN1, and PSD-95 from cultured hippocampal neurons infected on DIV7 with AAV-EGFP or AAV-CYLD and lysed on DIV21. (B) Quantifications of western blot band intensity from (A) relative to β-actin. n = 4 (WT) and 4 (KO). * $p < 0.05$; two-tailed unpaired t-test.

**Figure S4. Cortex autophagy blots. (**A) Representative western blots of LC3-I and -II from CYLD WT and KO cortical tissues. (B) Quantifications of LC3-I, LC3-II, and LC3-II/I ratio from (A), relative to β-actin. (C) Representative western blots of total and phosphorylated Akt and mTOR from CYLD WT and KO cortical tissues. (D) Quantifications of total and phosphorylated Akt and mTOR from (C), relative to β-actin. n = 6 (WT) and 6 (KO). * $p < 0.05$, ** $p < 0.01$; two-tailed unpaired t-test.

**Figure S5. Rapamycin enhancement of autophagy in the brain and rescue of synaptic proteins and mEPSCs.** (A) Representative western blots of GluA1, LC3-I and LC3-II of hippocampal lysates from P29-30 CYLD WT and KO mice treated with 1.5 mg/kg/day rapamycin or saline from P20-P28. (B) Quantifications of (A), relative to β-actin. (C) Cumulative probability plots of mEPSC interevent interval (C) and amplitude (D) of CA1 hippocampal neurons from treated and untreated WT and KO mice. ## $p < 0.01$, ### $p < 0.001$ Kolmogorov Smirnov test vs. WT + Saline.

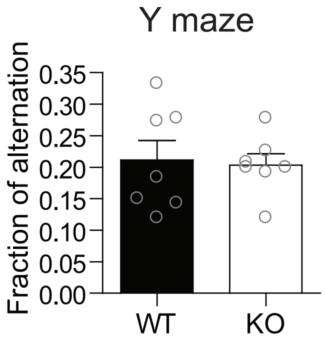
Y maze
Fraction of alternation
0.35
0.30
0.25
0.20
0.15
0.10
0.05
0.00
WT
KO

Figure S1

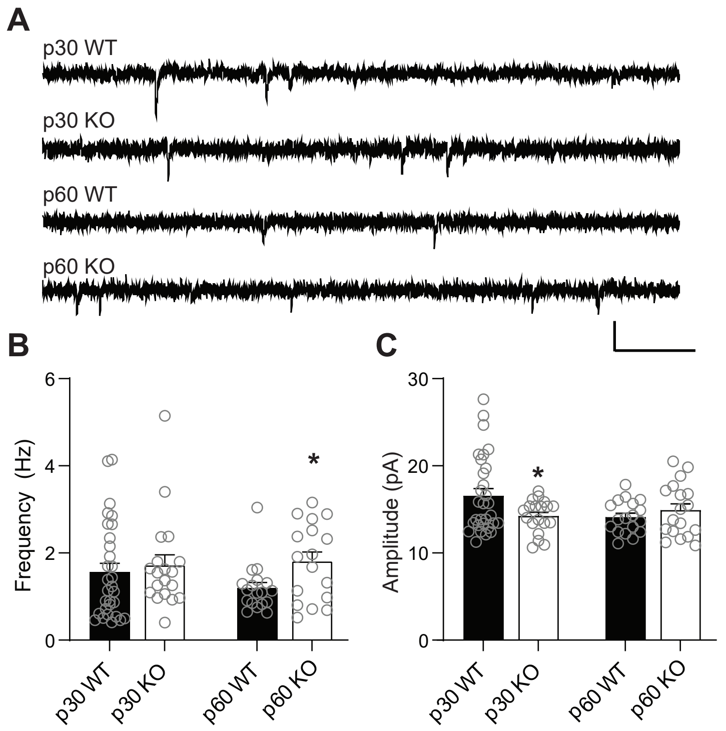
A
p30 WT
p30 KO
p60 WT
p60 KO
B
Frequency (Hz)
6
4
2
0
*
p30 WT
p30 KO
p60 WT
p60 KO
C
Amplitude (pA)
30
20
10
0
*
p30 WT
p30 KO
p60 WT
p60 KO

Figure S2

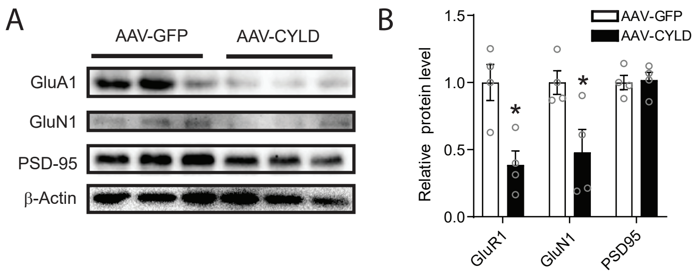
A
AAV-GFP
AAV-CYLD
GluA1
GluN1
PSD-95
β-Actin
B
AAV-GFP
AAV-CYLD
Relative protein level
1.5
1.0
0.5
0.0
*
*
GluR1
GluN1
PSD95

Figure S3

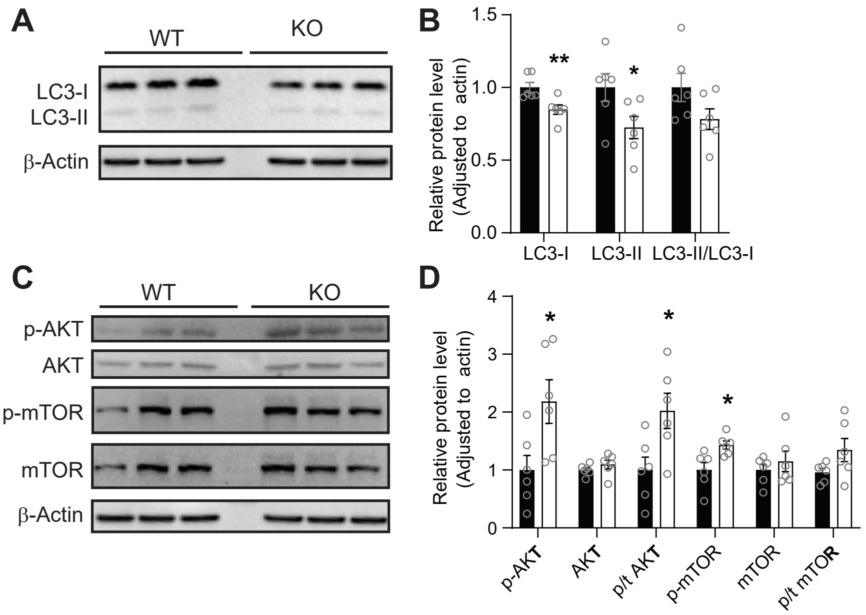
A
WT
KO
LC3-I
LC3-II
β-Actin
B
Relative protein level (Adjusted to actin)
1.5
1.0
0.5
0.0
**
*
LC3-I
LC3-II
LC3-II/LC3-I
C
WT
KO
p-AKT
AKT
p-mTOR
mTOR
β-Actin
D
Relative protein level (Adjusted to actin)
4
3
2
1
0
*
*
*
p-AKT
AKT
p/t AKT
p-mTOR
mTOR
p/t mTOR

Figure S4

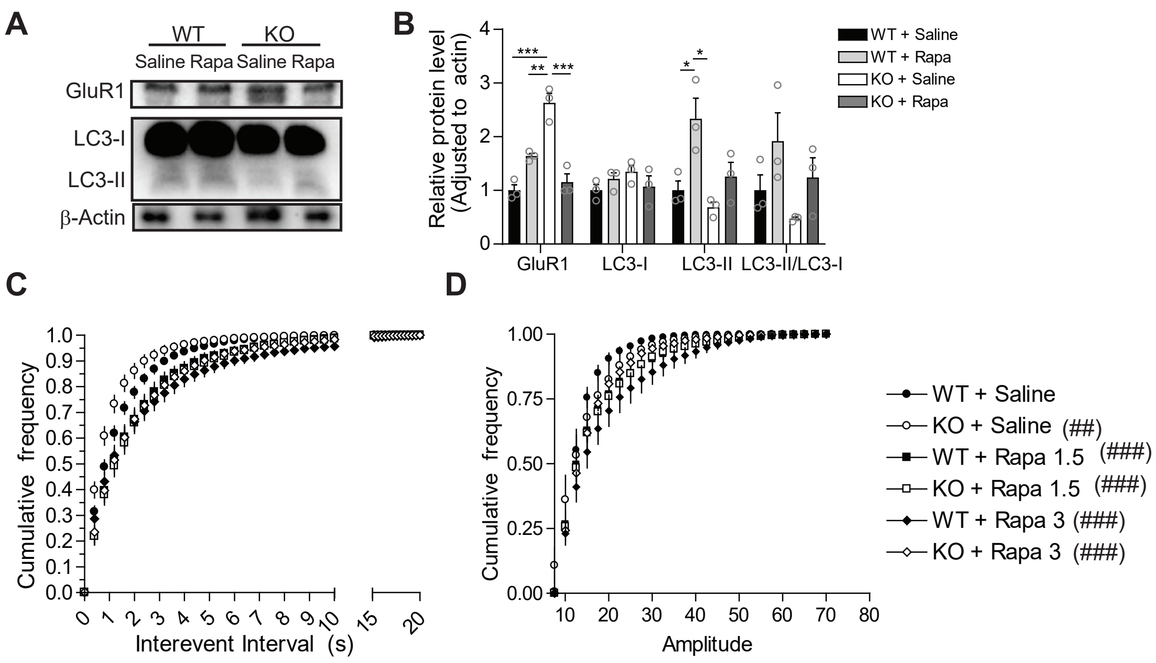
A
WT
KO
Saline Rapa Saline Rapa
GluR1
LC3-I
LC3-II
β-Actin
B
Relative protein level (Adjusted to actin)
WT + Saline
WT + Rapa
KO + Saline
KO + Rapa
GluR1
LC3-I
LC3-II
LC3-II/LC3-I
C
Cumulative frequency
Interevent Interval (s)
D
Cumulative frequency
Amplitude
WT + Saline
KO + Saline (##)
WT + Rapa 1.5 (###)
KO + Rapa 1.5 (###)
WT + Rapa 3 (###)
KO + Rapa 3 (###)

Figure S5
